# An Automated Behavioral Apparatus to Assess Distal Forelimb Function in Non-Human Primates

**DOI:** 10.1101/396572

**Authors:** Douglas Totten, Lisa Novik, Kari Christe, Marie-Josee Lemoy, Jeffrey Roberts, Jose Carmena, Robert Morecraft, Karunesh Ganguly

## Abstract

**Background:** Primate distal forelimb behaviors are commonly assessed using reach-to-grasp tasks. While these tasks are widely recognized as sensitive assays for forelimb function, they often require experimenter input, lack precise temporal cues for physiological monitoring, and can be expensive.

**New Method:** Using components developed by open-source electronics platforms, we have designed and tested a low-cost system to measure distal forelimb function in non-human primates. Our system is inexpensive; it is made primarily of acrylic and 3D printed plastic parts. Our control software, developed in MATLAB, was also used to control two cameras in order to capture and process video during behavior. The system was equipped with sensors, motors and microcontrollers to control the timing of the task and facilitate synchronization between behavior and neurophysiology with high temporal precision.

**Results:** We demonstrate that this system can be used to monitor motor recovery after stroke and investigate neurophysiological correlates of motor control.

**Comparison with Existing Methods:** Compared to a previous version of this task, our setup reduces experimenter input while providing unbiased delivery of behavioral cues and behavioral measurements with the temporal precision necessary for electrophysiological studies.

**Conclusions:** In summary, our system will allow unbiased monitoring of forelimb function in both healthy and injured animals that is compatible with electrophysiological studies.

## Introduction

There is growing interest in understanding the neurophysiological correlates of dexterous actions in non-human primates (Schaffelhofer and Scherberger, 2016; Vaidya et al., 2015). Traditionally, neurophysiological investigations of primary motor cortex (M1) have focused on the neural correlates of gross reaching movements, e.g. point-to-point reaches with tracking of the hand as a kinematic ‘end-point’ (Chestek et al., 2007; Ganguly and Carmena, 2009; Georgopoulos et al., 1986, 1988; Lee et al., 1998). Such research has resulted in important descriptions of both single neuron and population responses during ‘center-out’ reach tasks (Churchland et al., 2012; Georgopoulos et al., 1986, 1988; Lee et al., 1998). However, it is also readily clear that an important function of M1 is for dexterous hand control (Darling et al., 2009; Hoffman and Strick, 1995; Murata et al., 2008). This is particularly clear when examining recovery of forelimb function after a M1 lesion; while proximal functions recover quickly, recovery of distal hand function often takes significantly longer (Darling et al., 2011; Ganguly et al., 2013; Lawrence and Kuypers, 1968). Moreover, while there are automated tasks for assessing distal hand function they are not readily compatible with neurophysiological investigations (Darling et al., 2006; Gash et al., 1999; Pizzimenti et al., 2007). Our primary goal was to develop and validate a low-cost, automated system to allow neurophysiological monitoring during the performance of both reaching and grasping tasks. We particularly focused on recovery after a M1 lesion in order to demonstrate its utility for automated assessment of motor recovery.

Traditional approaches to assessing distal motor function require extensive experimenter input and do not provide precise temporal cues amenable for neurophysiological monitoring (Nudo and Milliken, 1996; Friel et al., 2005; Pizzimenti et al., 2007; Schmidlin et al., 2011; Gash et al., 1999, p. 199). Such tasks commonly involve “Kluver boards” or “Brinkman boards” that require dexterous manipulation (Nudo and Milliken, 1996; Friel et al., 2005; Pizzimenti et al., 2007; Schmidlin et al., 2011). Targets are presented in a fixed-location or limited area to allow researchers to investigate distal motor control. By comparison center-out tasks involve presenting similar targets at separate locations in space and primarily require reaching movements without a grasp component (Chestek et al., 2007; Georgopoulos et al., 1986, 1988; Lee et al., 1998). Our specific focus was to minimize the need for user input and supervision during the training and assessment of animals. Here we present two experimental devices constructed from inexpensive motors and microcontrollers, and parts that can be manufactured with a high degree of reproducibility using 3D printing and laser cutting.

The first task is a fully-automated reach-to-grasp task (RTG) where subjects retrieve assorted food-pellets from plastic wells of varying sizes; well size can be varied parametrically to control task difficulty. The second task is a center-out style reach-to-grasp (RTG_CO_) task where subjects retrieve small food-rewards from evenly spaced windows surrounding a central initiation sensor. Both tasks involve holding an initiation sensor for a short hold period before being presented with the food reward. Trial initiation and reward presentation were automated, thereby reducing the need for experimenter interventions. Further, the use of an open-source microcontroller as well as off-the-shelf electronic sensors allows easy synchronization between reach and grasp behavior and neurophysiological data (Wong et al., 2015). Thus, our setups minimize experimenter input and include sensors that can be used to temporally decompose the neuron-behavior relationship. Importantly, the ability to automate assessments has the added benefit of facilitating blinding of assessments.

## 2. Methods

### 2.1 Subjects

Experimental apparatuses were tested on two male rhesus macaques between 5 and 7 years of age. Subjects had hands without any deficits. Subjects were food scheduled, performing tasks in the morning and not receiving access to food until after testing. Subjects were provided with species-appropriate environmental enrichment, fed chow twice daily (LabDiet Monkey Diet 5047, Purina Laboratory, St. Louis, MO, USA), offered water ad libitum via automatic watering devices, and supplemented with fruits and vegetables biweekly. Weights were monitored weekly throughout testing with supplemental feeding provided post stroke-induction surgery. The current study was approved by the IACUC of the University of California, Davis. Animals were maintained in accordance with the USDA Animal Welfare Act and regulations and the *Guide for the Care and Use of Laboratory Animals. (***Animal Welfare Act as Amended**. 2013. 7 USC §2131–2159. **Animal Welfare Regulations**. 2013. 9 CFR § 3.129. **nstitute for Laboratory Animal Research**. 2011. Guide for the care and use of laboratory animals, 8th ed. Washington (DC): National Academies Press.)The animal care and use program of the University of California, Davis is fully accredited by AAALAC-i, USDA-registered, and maintains a Public Health Services Assurance. (**National Institutes of Health**. 2002. Public health service policy on humane care and use of laboratory animals. Bethesda (MD): Office of Laboratory Animal Welfare.)

### 2.2 Experimental Apparatuses

During experimental testing, subjects were placed in a nonhuman primate chair (Custom, University of California National Primate Center, Davis, CA). The chair had a front door allowing subjects to interact with the space in front of the chair. Once this door was opened, a custom-built device platform was wheeled into place and attached to the chair. The platform had a thick vertical wall of high-density polyethylene, which covered the door opening. This wall had an aperture positioned so that one arm could extend from the chair and interact with the devices. Both devices were mounted on plastic sheets which could be attached onto the surface of the device platform using latches, allowing devices to be easily swapped. The mounting sheet for the RTG task positioned the task at a 10° angle from the bottom front edge of the device to accommodate visibility.

Acrylic structural components were cut from cast acrylic 0.56cm (or 1/4”) thick for the RTG task and 0.30cm (or 1/8”) and 1.12cm (or 1/2”) thick for the RTG_CO_ task (Tap Plastic, San Francisco, CA) using a laser cutter (VLS3.50 Universal Laser Systems, Scottsdale, AZ). Vector drawings for laser-cutter operation were drawn using Adobe Illustrator (version CC, Adobe, San Jose, CA). Stanchions, motor chassis, shelves, capacitive sensor screw-mount, wells, wheel-forks, and the pellet placement arm were 3D printed using acrylonitrile butadiene styrene and dissolvable support material (uPrint and uPrint-Plus, Stratasys, Eden Prairie, MN). A capacitive touch sensor (MPR121, Adafruit, New York City, NY) connected to a screw coated in copper tape served as the trial-initiation control. The capacitive touch sensor was hooked to an Arduino UNO microcontroller (Arduino Uno - R3, Arduino, Ivrea, Italy) loaded with the corresponding library (Adafruint_MPR121, Github.com). These Arduinos also controlled lights which served several functions, including trial-indicator lights that facilitated offline clipping of videos into individual trials, lights which indicated that the device was waiting for hold-initiation, and lights which indicate which shelf is the target (Supplemental Figure 2). The rotating components were driven by a high-torque stepper motor (17HS19-1684S-PG14, Stepperonline, Nanjing City, China), coupled with a screw hub (part number 545636, Servocity, Winfield, KS) and powered by stepper-motor driver (TB6600, Dfrobot.com, Shanghai, China) which was operated by a separate Arduino.

For the RTG task wells were 0.59cm deep, with a base diameter 0.5cm smaller than the opening diameter. Opening diameters tested here were 1.3cm, 1.9cm, 2.5cm, 3.1cm, and 3.7cm. A 1.0cm well was deemed too difficult and was not used. Wells had hinged bottoms allowing the pellets to be automatically cleared after the trial (Supplemental Figure 4). Pins for the well-hinges were made from 0.2cm thick stainless-steel rods (ASIN B00KHUR5AQ, Amazon.com).

For the RTG task, pellets were dispensed from an automated pellet dispenser (80209-190S, Lafayette Instrument, Lafayette, IN) connected to the pellet-dispenser arm using a short strip of surgical tubing with an inner diameter of 3/8” (0.95cm). For the RTG_CO_ task, small food rewards were placed on the shelves (2.0cm wide by 2.25cm deep by 0.75cm tall) while access to the shelf was blocked. If the reward was not retrieved before the trial ended, it was removed manually after the trial.

The capacitive-touch sensors were connected to wires which were crimped onto screws, which served as the trial-initiation sensor. For the RTG_CO_ task, this screw was held in place by a 3D printed screw mount which allowed a non-conductive coupling to the screw hub (Supplemental figure 8).

### 2.3 Computer control of devices

Arduinos communicated with MATLAB (R2016b) via custom functions/Arduino sketches that communicated via a serial connection. Functions and sketches are available upon request.

### 2.4 Behavioral monitoring through synchronized video recordings

Two cameras (CM3-U3-13Y3C-CS, Point Grey, Richmond, BC, Canada) were mounted on a stainless-steel frame attached to the device platform. Camera shutters were controlled via Arduino-generated 5V trigger pulses delivered through the general-purpose input/output (GPIO) ports, allowing synchronous image acquisition. These trigger-pulses were recorded by the electrophysiology rig, thus allowing for camera/physiology synchronization. For each trial capacitive-sensor release, reach-start, grasp-start, grasp finish, number of attempts, and a trial-score were assigned.

Capacitive-sensor release was automatically detected from the sensor state which was recorded by the electrophysiology rig; reach-start was defined as the moment the center of mass of the hand began moving towards the target; grasp start was defined as the moment a finger came in contact with the well (RTG) or broke the plane of the reward window (RTG_CO_); grasp finish was the moment the pellet was securely gripped and began retracting (RTG) or the reward fully passed the plane of the reward window (RTG_CO_); number of attempts was the number of contiguous periods of contact with the reward; trial scores where either 1: reward successfully retrieved, 2: reward removed from well/shelf but not maintained in a grasp (e.g. dropped), 3: reward not successfully removed from well/shelf, 4: invalid trial (i.e. subjects did not attempt to retrieve a reward). Grasp duration was calculated as time from grasp start to grasp finish. Percent success was calculated as

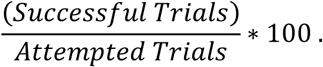

Cameras were calibrated using synchronous images of checkerboards and MATLAB’s Stereo Camera Calibration app. Each pair of synchronous annotations were used to triangulate the finger location using the camera calibration matrices (see below).

### 2.5 Behavior training

Once subjects adapted to the primate chair and received rewards placed in the devices, they began executing trials with the device lights/motors active. Trial-counts were gradually increased over training days. Subjects were trained for approximately one month on the final task design prior to stroke-induction, at which point they had stable behavioral scores over three days and could perform a minimum of 50-trials/day for each task. To maintain a high-level of motivation, subjects received bonus rewards after successfully completing several trials which could be a liquid or solid food treat depending on subject preference.

Following stroke-induction, subjects were given a minimum 1-week postoperative recovery period prior to resuming testing. Upon resuming testing, subjects initially had difficulty consistently making contact with the initiation screw. As they recovered they were rewarded progressively until they were able to perform the task again. Each trial was initiated with a random hold-period between 0.5-0.8s, after which the target was presented by having the well rotate into an accessible position (RTG) or the window-blocker rotate out of the way (RTG_CO_).

### 2.6 Stroke induction and electrophysiology implantation surgery

Lesions and electrode implantation were performed in a single surgical session. Preoperatively, subjects were sedated with ketamine hydrocholoride (10 mg/kg), administered atropine sulfate (0.05 mg/kg), prepared and intubated. They were then placed on a mechanical ventilator and maintained on isoflurane inhalation (1.2-1.5%). Subjects were positioned in a stereotactic frame (David Kopf Instruments, Tujunga, CA) and administered mannitol (1.5 g/kg) intravenously prior to the craniotomy. A skin incision, bone flap, and dural flap were made over the lateral frontoparietal convexity of the hemisphere and the caudal region of the frontal lobe and rostral region of the parietal lobe was exposed unilaterally. After cortical exposure, the lesion was induced using surface vessel coagulation/occlusion followed by subpial aspiration targeting the forelimb region of primary motor cortex using anatomical landmarks. Following resection, a flap of the dura was sutured to cover the lesioned area, while leaving a small window anterior to the lesion for electrode implantation. A microwire recording array was targeted to the dorsal premotor cortex (PMd) using anatomical landmarks and inserted to approximately 2mm. Another microwire was implanted in the primary somatosensory cortex. The galea aponeurotica, temporalis muscle and skin were closed using standard surgical procedures. Each animal was carefully monitored post-operatively for 7 days.

### 2.7 Electrophysiology

Electrophysiological recordings were made using a ZIF-Clip based 64-channel microwire array, connected to a 128-channel PZ5, RZ2 processer, WS8 workstation and RS4 data streamer (Tucker-Davis Technologies, Alachua, FL). To extract spiking information all channels were referenced to a pair of titanium cranial screws in the contralateral hemisphere, bandpass filtered from 500-10,000hz, processed at 24414hz and thresholded to detect events exceeding 5-standard deviations. Waveforms were visually verified online.

One challenge we encountered using this setup was electrical noise interfering with the electrophysiological recordings. In particular, there was potential for interference while subjects were touching the capacitive touch sensors. A large portion of the electrical noise was mitigated by ensuring common grounding of all Arduino boards. There were further complications with the capacitive-touch sensor in the RTG_CO_ task where the sensor is part of the same physical assembly as the motor drive shaft. This was addressed by building a 3D printed plastic mount for the screw that served as the capacitive touch sensor. This isolated the sensor from the conductive set-screw hubs (Supplemental Fig. 7).

### 2.8 Statistical analysis

To calculate reach-related tuning, firing rates in the period 500ms prior to capacitive-touch sensor release were compared to the target angle using established methods. Briefly, the firing rate was estimated from the target angle using the following equation

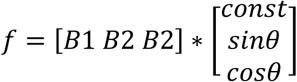

Where *f* represents the firing rate, θ represents the target angle. The coefficients, *B*, were estimated using linear regression (Ganguly and Carmena, 2009). A unit had significant reach-direction tuning if the linear regression had a p<0.05. Preferred direction was calculated as tan^-^ 1(B_2_/B_3_). Modulation depth was the difference between the maximum and the minimum of the function *f.*

Peri-event time histograms were constructed by aligning all the trials on a day to capacitive-touch sensor release or grasp-start, then using a 200ms bin slid in 25ms increments. The bins with starting times from 2s to 1.5s prior to the event were used to generate a baseline firing distribution. Each bin was determined significantly different from this baseline distribution if they had a p-value of less than 0.05 using a Wilcoxon rank-sum test.

RTG_CO_ reach trajectories were reconstructed from stereo-camera video for a subset of the data. This required manually annotating the thumb and forefinger locations in each video stream. PCA was used to transform the 3D finger and thumb location estimates, and the first two components were found to approximately align with the vertical plane of the device revealing distinct, stereotyped reach trajectories to each target. Visual occlusion, precision in the manual annotation, and calibration accuracy limit the capabilities of such kinematic reconstruction.

## 3. Results

### 3.1 Behavioral performance in well-trained subjects

Performance during the RTG task was assessed by measuring the grasp-duration and number of attempts for successful trials, as well as the percentage of successful trials. All subjects were trained for >1 month to bring them to a performance plateau prior to stroke-induction. Fig. 2 shows behavioral performance for MMU127 and MMU986 averaged over the last 3 training sessions prior to stroke induction, separated by well size. Grasp duration was longest, number of attempts highest and percent success lowest for the 13mm well, with performance improving as the well size increased.

**Figure 1.**
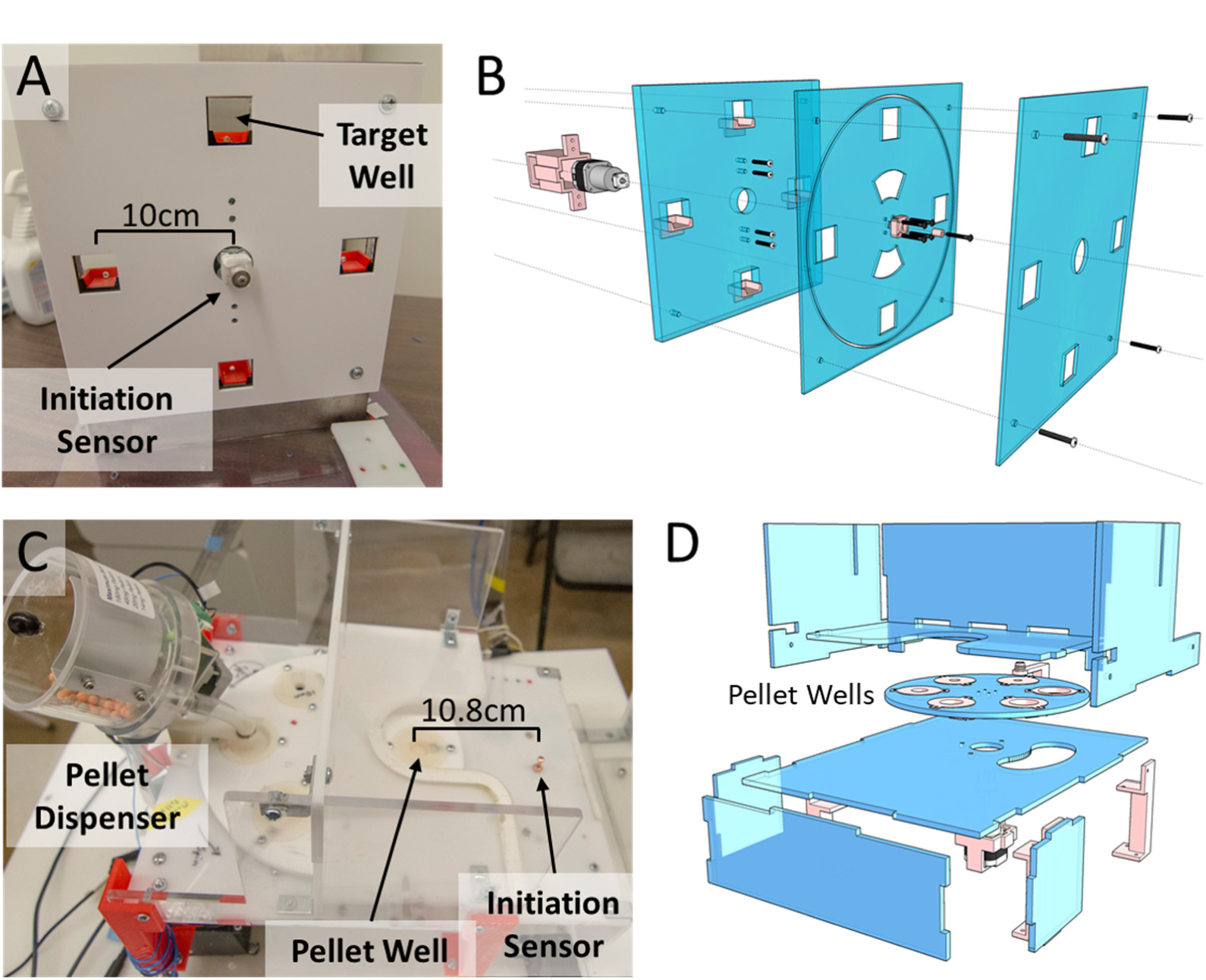
Description of devices. (A, B) Respective reach-to-grasp **‘**center-out’ (RTG_CO_) and (C,D) reach-to-grasp (RTG) photo and schematic.

**Figure 2.**
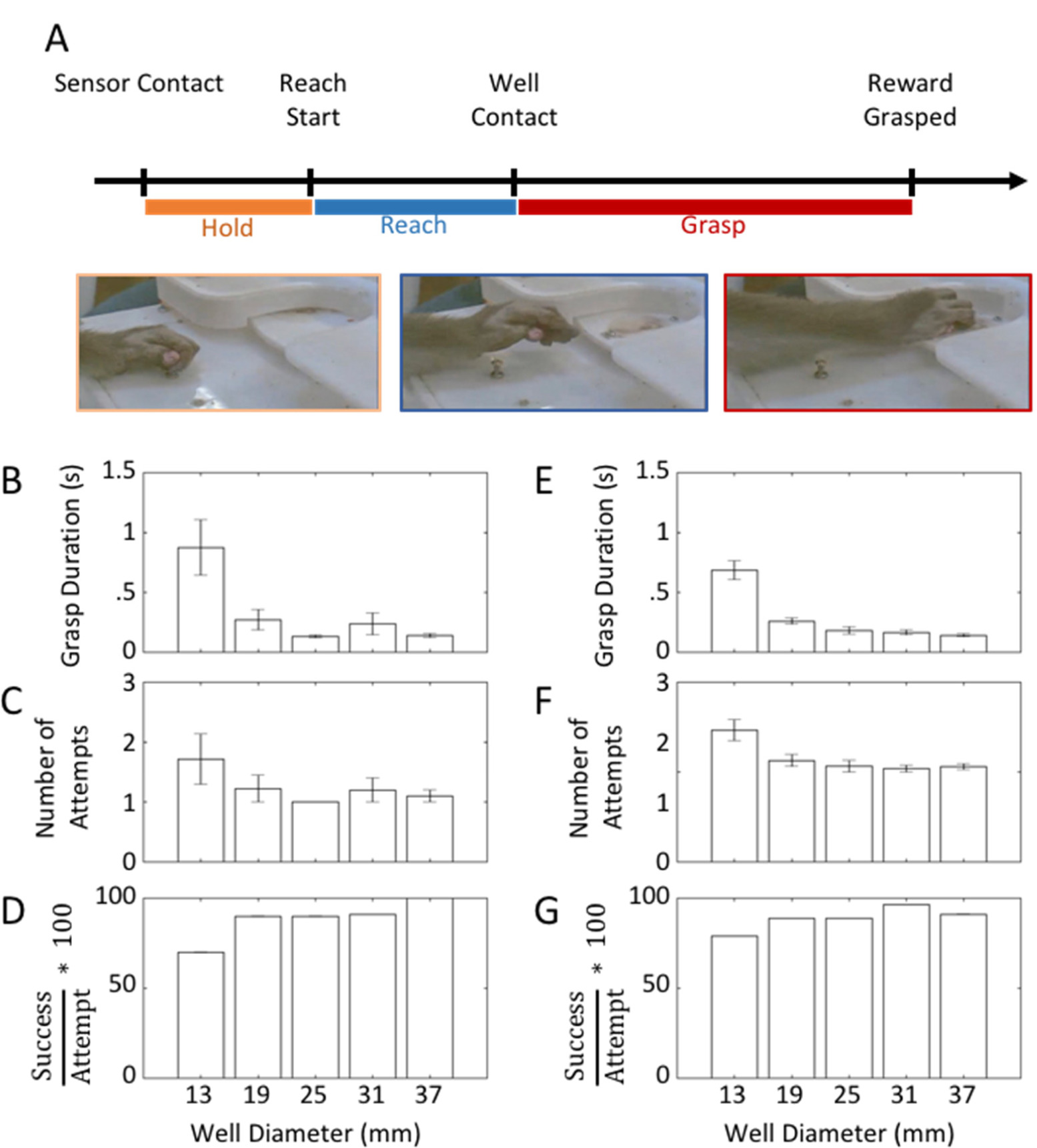
Behavioral performance. (A) Task design. Images from the RTG task are shown at respective points. Grasp-duration (B), number of attempts (C), and percent success (D) during the RTG task averaged across 11-presentations of each well for one representative test session from case MMU127. Error bars indicate standard error. (E), (F) and (G), same conventions as (B), (C), and (D) for three days for MMU127 and MMU986, for a total of 58 presentations of each well.

### 3.2 Fully-Automated Reach-to-grasp

Following a unilateral lesion to the forelimb region of the primary motor cortex, MMU127 and MMU986 demonstrated clear performance deficits in the RTG task that recovered with time (Fig. 3). Both subjects demonstrated decreased grasp durations that eventually recovered to pre-stroke performance levels (Fig. 3A,C). MMU986 showed a persistent deficit in percent success (Fig. 3B), while MMU127 demonstrated recovery with both metrics (Fig. 3C, D).

**Figure 3.**
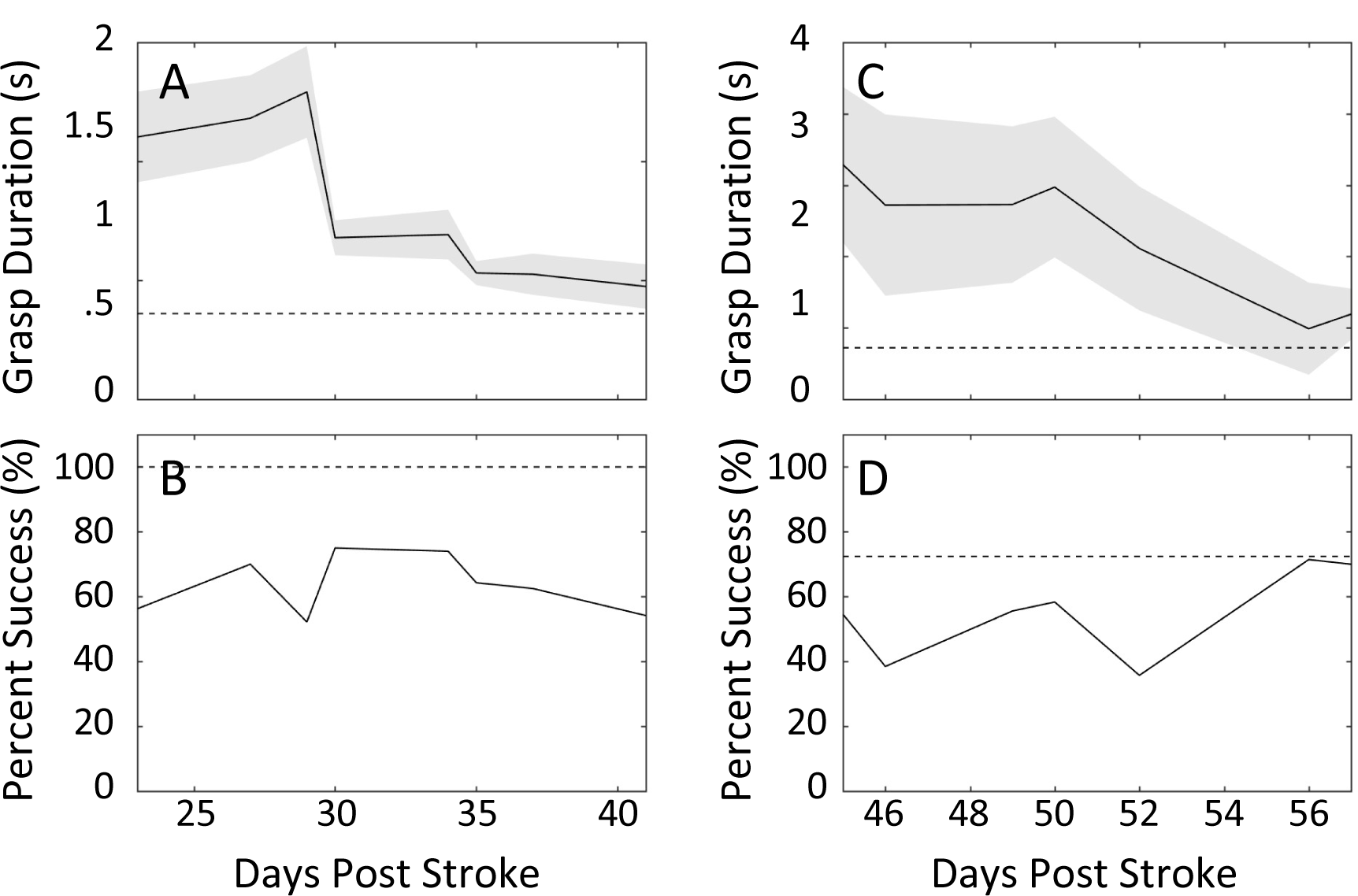
Reach-to-grasp performance recovery. Grasp duration (A) and percent success (B) for MMU986. Solid line shows the average performance and shading indicates the standard error. Dashed line shows pre-stroke performance, averaged over 3 sessions. Grasp duration (C) and percent success (D) for MMU127.

Here we analyzed data from the 64-channel microwire array implanted in PMd, which was in the anterior perilesional area. Neural responses were time-locked to capacitive touch sensor release, or behavioral timepoints identified from videos including reach start, grasp start and grasp finish (Fig. 4A). Figure 4B and C show responses from MMU986 unit35 for 25-trials retrieving pellets from a 1.3cm well with responses locked to capacitive touch-sensor release, exhibiting clear event-related firing rate modulation. This was investigated across units by calculating the number of channels with significant increases or decreases from baseline firing over time relative to capacitive-touch release (Fig. 4D). The same can be seen for responses locked to grasp-start for MMU986 unit60 (Fig. 4E, F) and across the population (Fig. 4G).

**Figure 4.**
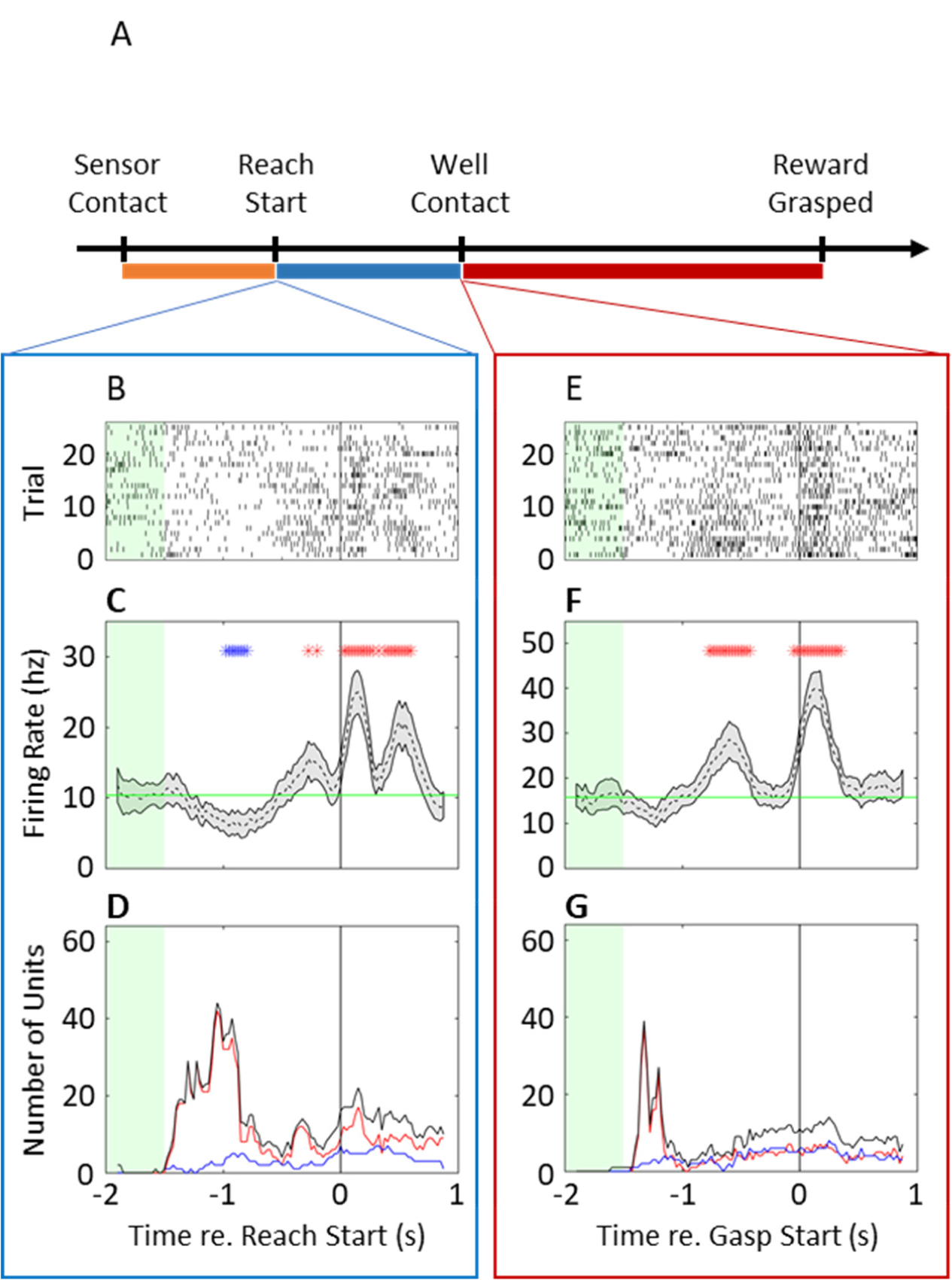
Neurophysiology during RTG task. (A) T Raster (B) and peri-event time histogram (C) for an example unit with responses aligned to the release of the capacitive touch sensor. The baseline period is indicated by the green band, with significant increases from baseline indicated by red stars, and significant decreases indicated by blue stars. (D) Number of units with firing rates significantly deviating from baseline at each time point when aligned to release of the capacitive touch sensor. (E), (F), and (G) have the same conventions as (B-D), except for a different unit an aligned to grasp-initiation.

### 3.3 Center-Out Style Reach-to-Grasp

It is common to observe neurons in the PMd with pre-reach firing that significantly varies with reach direction (Cisek and Kalaska, 2005; Takahashi et al., 2017). Peri-event time histograms and raster plots (Fig. 5A) aligned to capacitive-sensor release showed activity that was significantly modulated by target location. One representative testing day for MMU703 showed significant reach-related activity for 21 out of 64 units in the 500ms before capacitive-sensor release (Fig. 5B).

**Figure 5.**
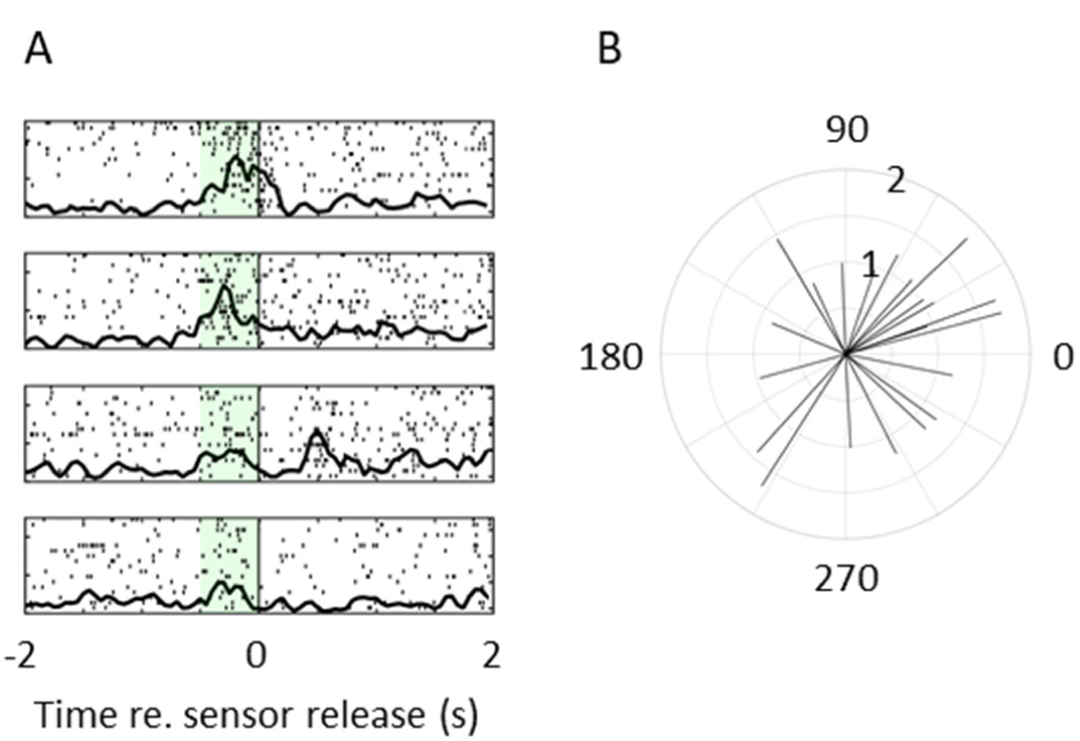
Neurophysiology during the RTG_CO_ task. (A) Peri-event time histogram and raster for an example unit with responses aligned to the release of the capacitive touch sensor. Responses are separated by reaches towards targets at 0°, 90°, 180° and 270°, from top to bottom. The green band indicates the time-period used to detect significant reach-direction tuning. (B) Population vector for 21 units with significant reach-direction preference. The angle of the line indicates the preferred direction, while the length indicates modulation depth. Data for all figures is from one day of testing from MMU703.

## Discussion

In summary, our results demonstrate that our automated forelimb assessment apparatus can facilitate initial training and assessment of motor recovery after injury as well as be compatible with neurophysiological monitoring. The incorporation of wait and trial cues allowed us to easily lock neurophysiological data to relevant parameters. The temporal resolution was such that it could be easily integrated with our electrophysiological workstation.

An advantageous feature of our design is that the wells are modular and easy to swap in and out, thus allowing further customization of the task. Other research has made use of modified reach-to-grasp designs that use slots or crosses to constrain the grasping behavior (Schmidlin et al., 2011). Such alterations could be easily implemented while retaining automation and experimenter/subject isolation. Given its flexibility, the setup is also amenable to evaluating a wide spectrum of models; well ports are designed to accommodate a range of possible well-sizes and as well as non-well elements. For example, it is possible to change the well for novel objects for learning and/or studies of dexterous manipulation (Schaffelhofer and Scherberger, 2016).

While this setup has significantly improved our ability to train and monitor behavior, there are several remaining cumbersome steps. This includes assessment of kinematics and behavioral scoring (e.g. duration of grasping). This still requires manual scoring; albeit we have developed graphical user interfaces that can improve this process. Future areas of development include the addition of either marker based (Rouse and Schieber, 2015) or markerless tracking (Overduin et al., 2010; Schaffelhofer and Scherberger, 2012). Moreover, the addition of sensors to the objects themselves can allow us to determine the moment of contact as well as other trial parameters.

It should be noted that while reach-to-grasp and center-out tasks do an excellent job of dissociating distal and proximal motor control, the dynamic coupling of motor systems makes complete isolation impossible (Rouse and Schieber, 2016, 2015; Saleh et al., 2012). It is reasonable to expect that wells of different sizes could lead subjects to supinate/pronate in preparation for different grasp strategies, and that retrieving rewards with the arm at different angles will necessitate different hand postures. While this is true, the features of interest in these tasks, namely grasp duration, number of attempts, and success rate for reach-to-grasp, or reach-trajectories and reach-direction tuning for the center-out task are independent of this kinematic detail. Future studies will focus on refining kinematic assessments to further assess such aspects of object manipulation.

## Conclusion

The RTG and RTG_CO_ are widely used methods for examining distal and proximal forelimb motor control in non-human primates respectively. There are several issues limiting the utility of these setups in research, including a trade-off between the cost of commercial solutions versus reproducibility from custom-solutions, the ease and reliability for training non-human primates to use them, the requirement for experimenters to interact with the subjects and challenges of associating neural activity with behavior. Here we present inexpensive customized setups that incorporate capacitive touch sensors for trial initiation allowing easy synchronization between behavior and neurophysiology. We are hopeful that this will allow greater translational approaches to understanding the network basis of recovery and neural engineering based treatments (Gulati et al., 2015; Ramanathan et al., 2018).

## Acknowledgements

This research was funded via a pilot grant from the California National Primate Research Center (CNPRC, Pilot Award #8777) and a Career Award for Medical Scientists from the Burroughs Wellcome Fund to K.G.

## Tables

**Table 1.** RTG parts list.

| RTG |  |  |  |  |  |  |
| --- | --- | --- | --- | --- | --- | --- |
| Part | Size | Distributer | Part Number | Quantity | Cost per Part | Total Cost |
| Acrylic | 0.56 x 30.48 x 60.96 cm | Amazon |  | 4 | 16.8 | 67.2 |
| Stepper Motor | Nema 17 | Stepperonline.com | 17HS19-1684S-PG14 | 1 | 36.1 | 36.1 |
| Motor Hub | 8mm bore | Servocity.com | 545636 | 1 | 4.99 | 4.99 |
| Motor Driver |  | dfrobot.com | TB6600 | 1 | 20 | 20 |
| Capacitive touch sensor |  | Adafruit.com | MPR121 | 1 | 12.5 | 12.5 |
| Pellet Dispenser |  | Lafayette Instruments | 80209-190S | 1 | 795 | 795 |
| Arduino UNO R3 |  | Arduino | A000066 | 2 | 22 | 44 |
| Steel Rods | 0.2cm dia. | Amazon | ASIN B00KHUR5AQ | 1 | 6.9 | 6.9 |
| Total |  |  |  |  |  | 986.69 |

**Table 2.** RTG_CO_ parts list.

| Part | Size | Distributor | Part Number | Quantity | Cost per Part | Total Cost |
| --- | --- | --- | --- | --- | --- | --- |
| Acrylic | 0.3 x 30.48 x 60.96 cm | Amazon |  | 1 |  | 0 |
| 1.12cm Acryluc | 1.12 x 30.48 x 30.48 cm | Amazon |  | 1 | 14.4 | 14.4 |
| Stepper Motor | Nema 17 | Stepperonline.com | 17HS19-1684S-PG14 | 1 | 36.1 | 36.1 |
| Motor Hub | 8mm bore | Servocity.com | 545636 | 1 | 4.99 | 4.99 |
| Motor Driver |  | dfrobot.com | TB6600 | 1 | 20 | 20 |
| Capacitive touch sensor |  | Adafruit.com | MPR121 | 1 | 12.5 | 12.5 |
| Arduino UNO R3 |  | Arduino | A000066 | 2 | 22 | 44 |
| Total |  |  |  |  |  | 131.99 |

## Supplemental Figures

**Supplemental figure 1.**
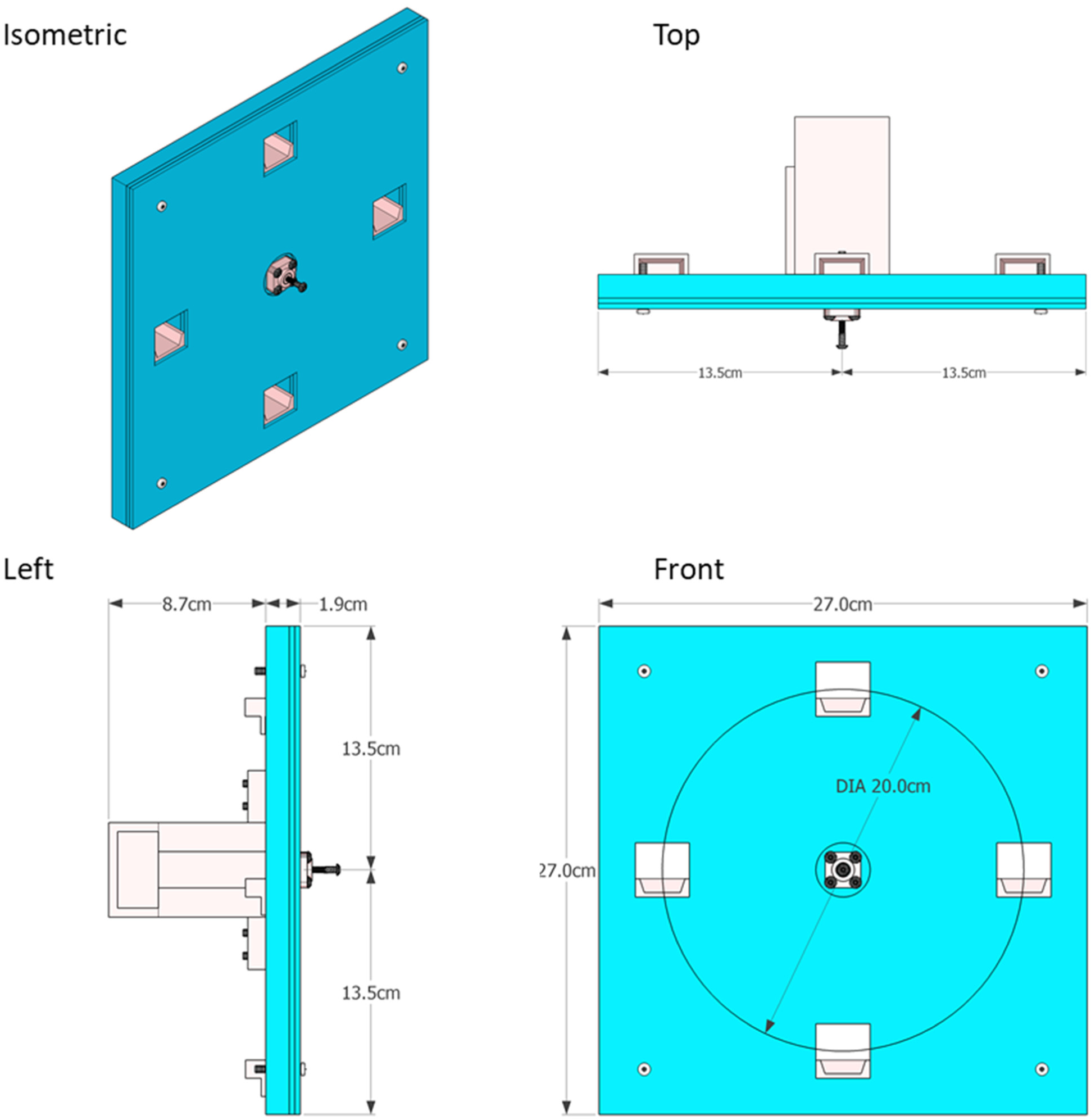
Assembled RTG_CO_ diagram with dimensions.

**Supplemental figure 2.**
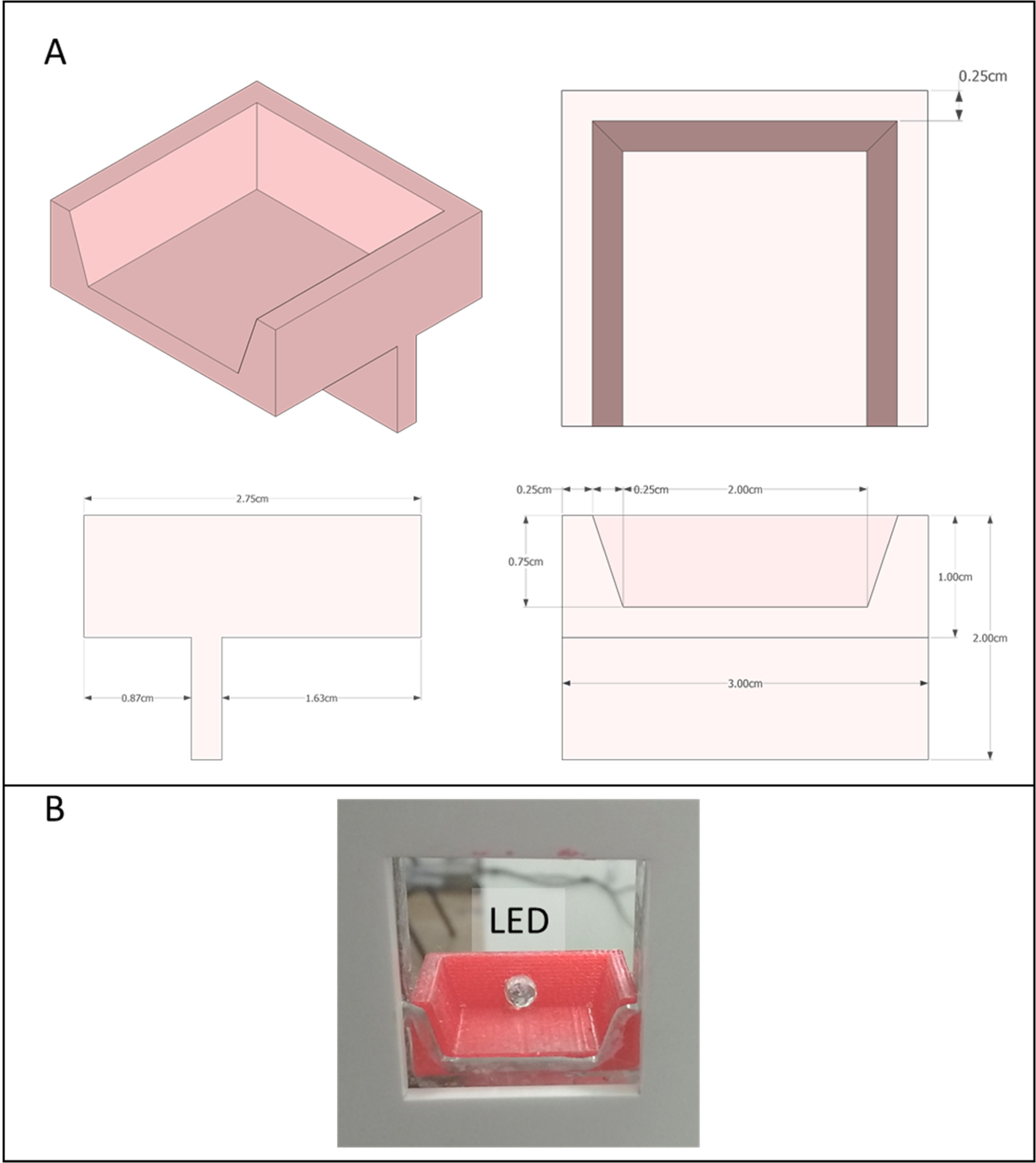
Design for the shelf for the RTG_CO_ task. *(A)* Dimensions. *(B)* Photo showing shelf from the front with LED added after printing.

**Supplemental figure 3.**
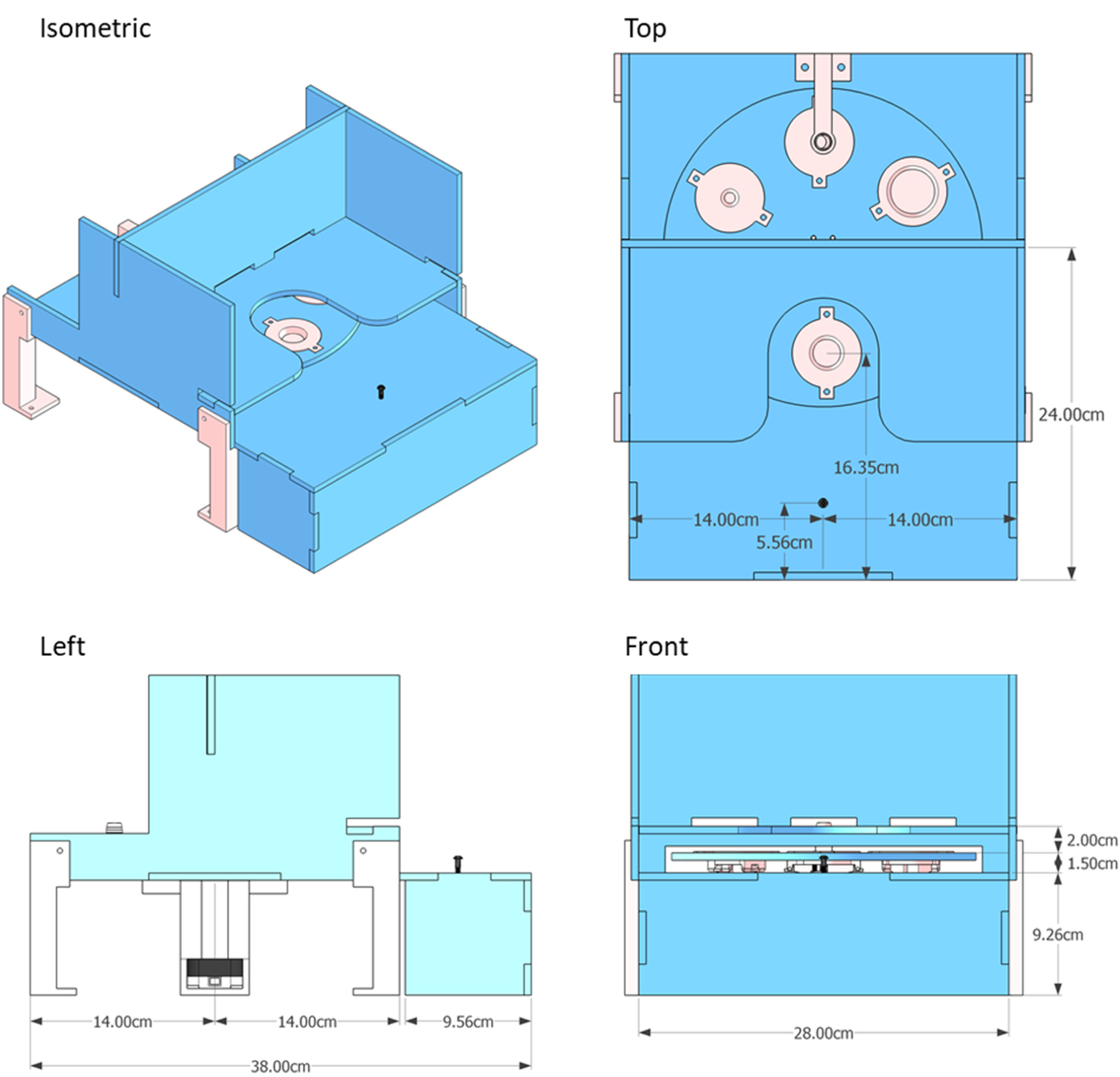
Assembled RTG diagram with dimensions.

**Supplemental figure 4.**
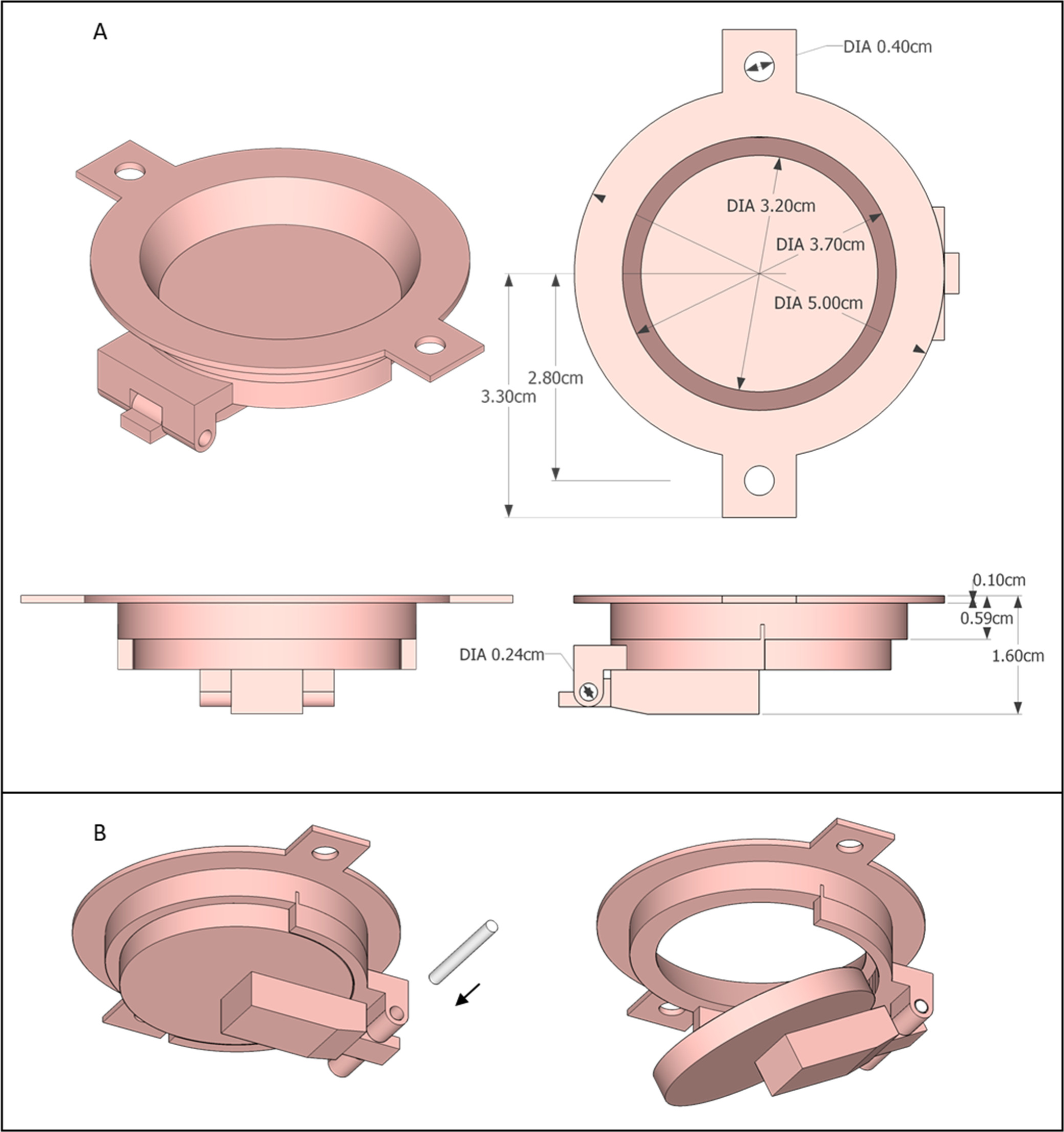
Well-insert dimensions details. (*A*) Isometric, top, side, and front views with dimensions. *(B)*, well assembly showing flap closed and pin insertion (left), and flap open with pin in place (right).

**Supplemental figure 5.**
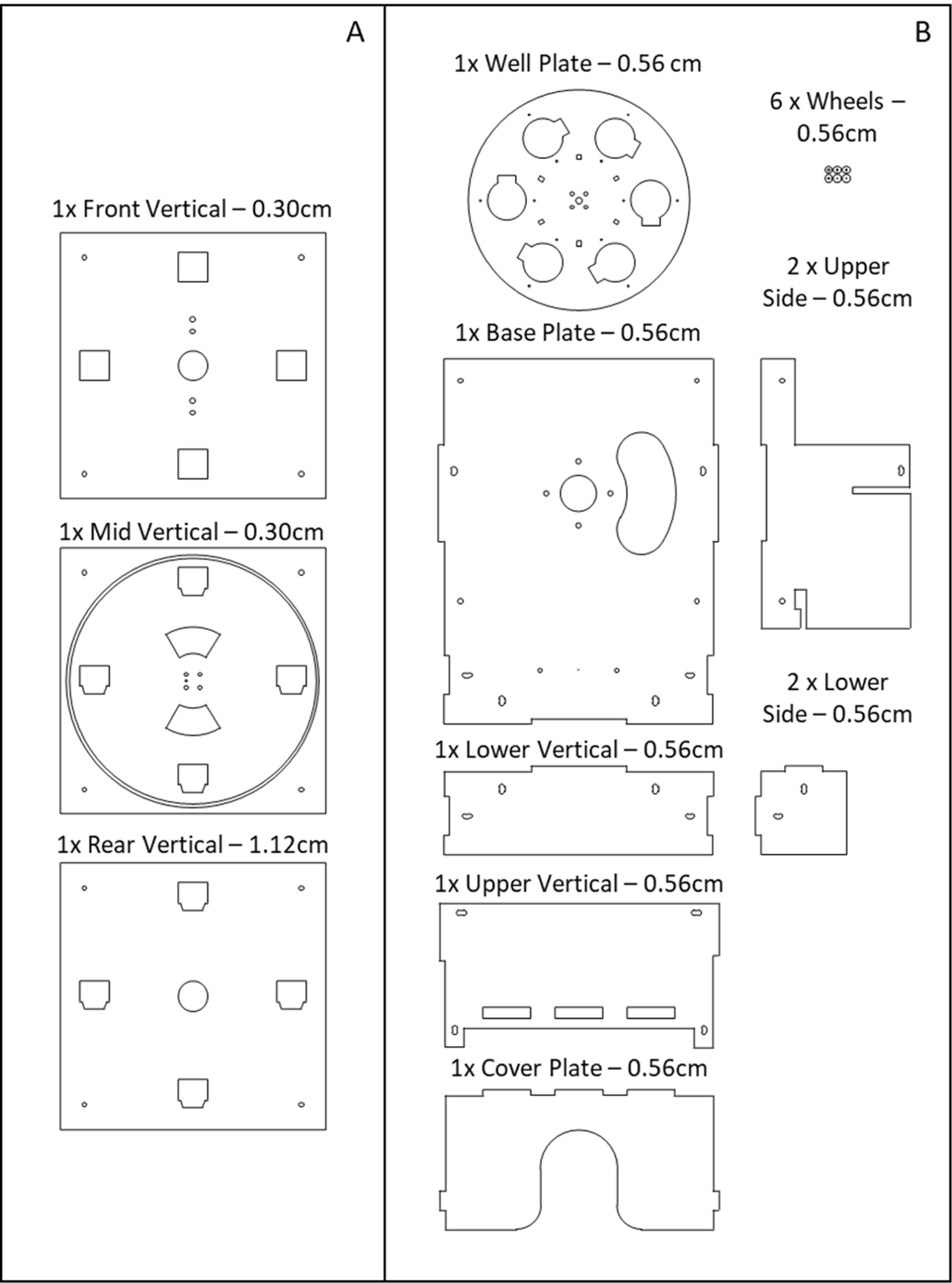
All laser-cut parts for RTG_CO_ (A), and reach to grasp (B) construction.

**Supplemental figure 6.**
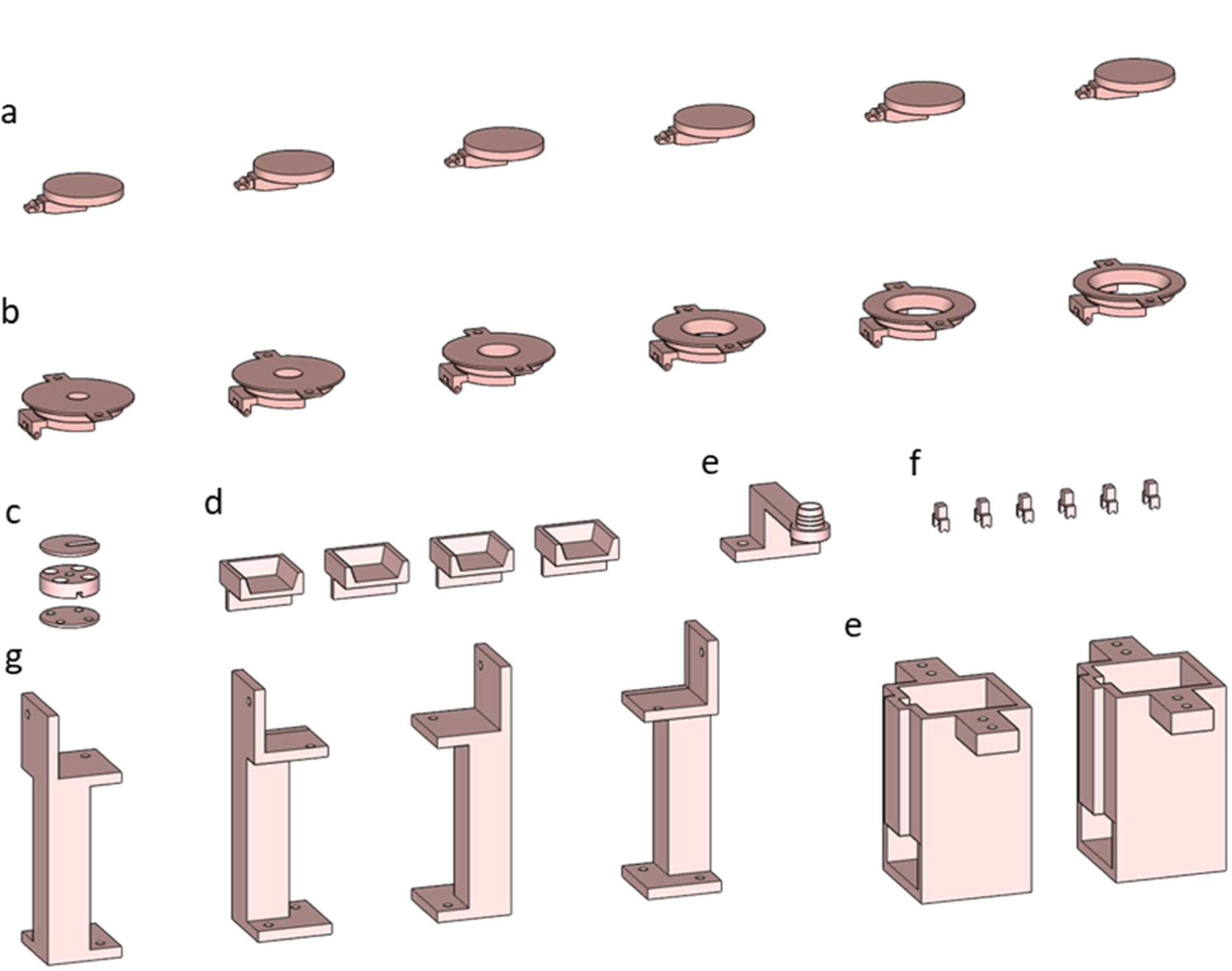
All 3D printed components. (a) 6x well flaps. (b) 6x wells. (c) 1x capacitive touch sensor screw mount. (d) 4x shelves. (e) 1x pellet placement arm. (f) 6x wheel forks. (g) 4x stanchions. (h) 2x motor chassis.

**Supplemental figure 7.**
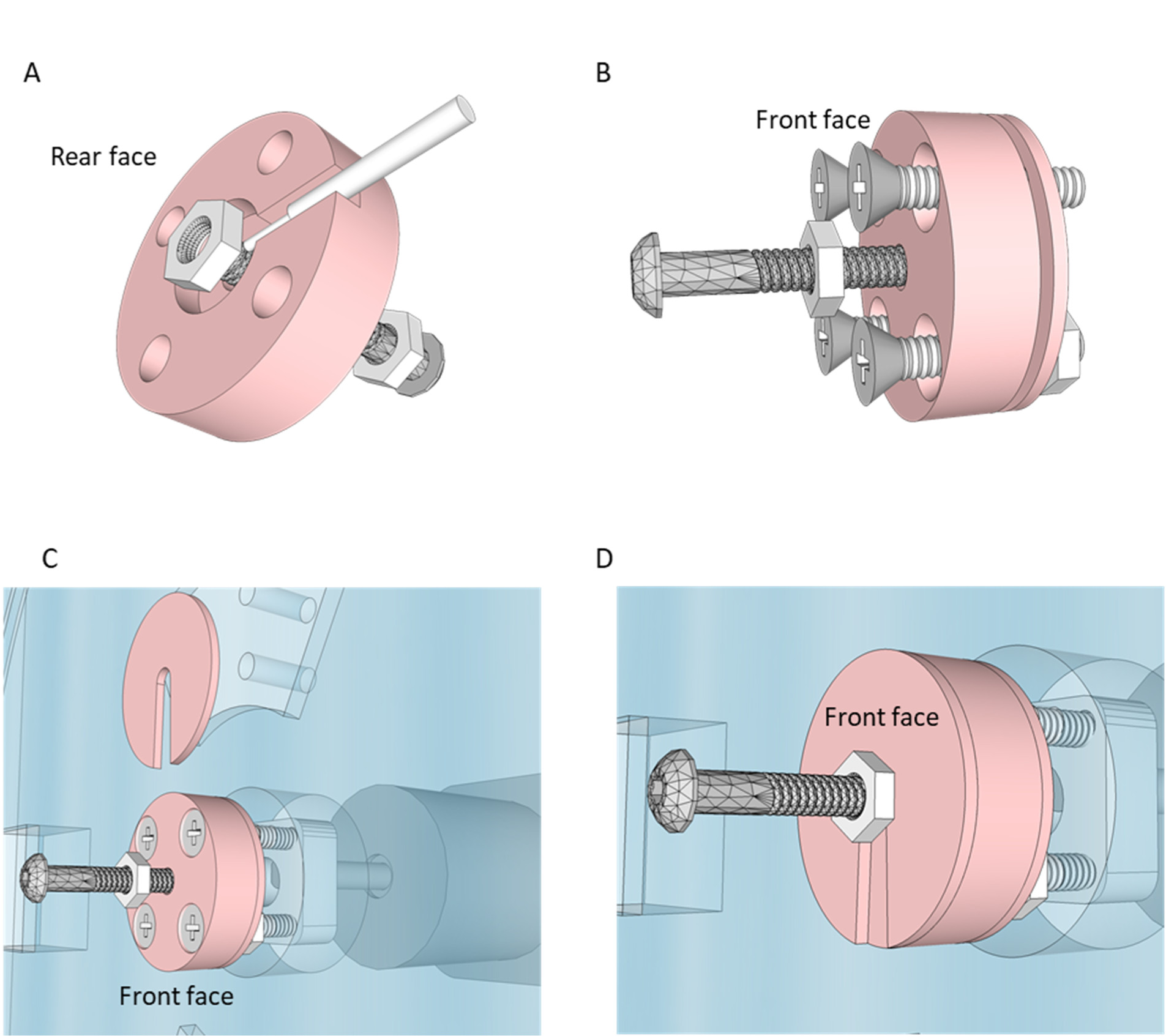
Capacitive sensor screw mount. For the RTG_CO_ task a screw in the center of the device serves as the capacitive touch sensor. A 3D printed plastic component is used to make a non-conductive mechanical attachment to the motor hub attached to the drive-shaft of the stepper motor. (A) The capacitive wire is crimped inside of the middle piece of the 3D printed part using a nut. Make sure a nut is on the central screw on what will be the front-face of the 3D printed part for attaching the front-shield (step *D*). (B) The rear-panel is slid in place isolating the capacitive wire. (C) The 3D printed component is screwed into the motor hub, with screws extending through the rotating component of the laser-cut acrylic. (D) The front shield is placed and held on with a nut.

## References

Chestek, C.A., Batista, A.P., Santhanam, G., Yu, B.M., Afshar, A., Cunningham, J.P., Gilja, V., Ryu, S.I., Churchland, M.M., Shenoy, K.V., 2007. Single-Neuron Stability during Repeated Reaching in Macaque Premotor Cortex. J. Neurosci. 27, 10742–10750. https://doi.org/10.1523/JNEUROSCI.0959-07.2007

Churchland, M.M., Cunningham, J.P., Kaufman, M.T., Foster, J.D., Nuyujukian, P., Ryu, S.I., Shenoy, K.V., 2012. Neural population dynamics during reaching. Nature. https://doi.org/10.1038/nature11129

Cisek, P., Kalaska, J.F., 2005. Neural Correlates of Reaching Decisions in Dorsal Premotor Cortex: Specification of Multiple Direction Choices and Final Selection of Action. Neuron 45, 801–814. https://doi.org/10.1016/j.neuron.2005.01.027

Darling, W.G., Peterson, C.R., Herrick, J.L., McNeal, D.W., Stilwell-Morecraft, K.S., Morecraft, R.J., 2006. Measurement of coordination of object manipulation in non-human primates. J. Neurosci. Methods 154, 38–44. https://doi.org/10.1016/j.jneumeth.2005.11.013

Darling, W.G., Pizzimenti, M.A., Morecraft, R.J., 2011. FUNCTIONAL RECOVERY FOLLOWING MOTOR CORTEX LESIONS IN NON-HUMAN PRIMATES: EXPERIMENTAL IMPLICATIONS FOR HUMAN STROKE PATIENTS. J. Integr. Neurosci. 10, 353–384. https://doi.org/10.1142/S0219635211002737

Darling, W.G., Pizzimenti, M.A., Rotella, D.L., Peterson, C.R., Hynes, S.M., Ge, J., Solon, K., McNeal, D.W., Stilwell-Morecraft, K.S., Morecraft, R.J., 2009. Volumetric effects of motor cortex injury on recovery of dexterous movements. Exp. Neurol. 220, 90–108. https://doi.org/10.1016/j.expneurol.2009.07.034

Friel, K.M., Barbay, S., Frost, S.B., Plautz, E.J., Hutchinson, D.M., Stowe, A.M., Dancause, N., Zoubina, E.V., Quaney, B.M., Nudo, R.J., 2005. Dissociation of Sensorimotor Deficits After Rostral Versus Caudal Lesions in the Primary Motor Cortex Hand Representation. J. Neurophysiol. 94, 1312–1324. https://doi.org/10.1152/jn.01251.2004

Ganguly, K., Byl, N.N., Abrams, G.M., 2013. Neurorehabilitation: motor recovery after stroke as an example. Ann. Neurol. 74, 373–381. https://doi.org/10.1002/ana.23994

Ganguly, K., Carmena, J.M., 2009. Emergence of a stable cortical map for neuroprosthetic control. PloS Biol. 7, 1508.

Gash, D.M., Zhang, Z., Umberger, G., Mahood, K., Smith, M., Smith, C., Gerhardt, G.A., 1999. An automated movement assessment panel for upper limb motor functions in rhesus monkeys and humans. J. Neurosci. Methods 89, 111–117.

Georgopoulos, A., Schwartz, A., Kettner, R., 1986. Neuronal population coding of movement direction. Science 233, 1416–1419. https://doi.org/10.1126/science.3749885

Georgopoulos, A.P., Kettner, R.E., Schwartz, A.B., 1988. Primate motor cortex and free arm movements to visual targets in three-dimensional space. II. Coding of the direction of movement by a neuronal population. J. Neurosci. 8, 2928–2937.

Gulati, T., Won, S.J., Ramanathan, D.S., Wong, C.C., Bodepudi, A., Swanson, R.A., Ganguly, K., 2015. Robust Neuroprosthetic Control from the Stroke Perilesional Cortex. J. Neurosci. 35, 8653–8661. https://doi.org/10.1523/JNEUROSCI.5007-14.2015

Hoffman, D.S., Strick, P.L., 1995. Effects of a primary motor cortex lesion on step-tracking movements of the wrist. J. Neurophysiol. 73, 891–895. https://doi.org/10.1152/jn.1995.73.2.891

Lawrence, D.G., Kuypers, H.G., 1968. The functional organization of the motor system in the monkey. I. The effects of bilateral pyramidal lesions. Brain J. Neurol. 91, 1–14.

Lee, D., Port, N.L., Kruse, W., Georgopoulos, A.P., 1998. Variability and Correlated Noise in the Discharge of Neurons in Motor and Parietal Areas of the Primate Cortex. J. Neurosci. 18, 1161–1170.

Murata, Y., Higo, N., Oishi, T., Yamashita, A., Matsuda, K., Hayashi, M., Yamane, S., 2008. Effects of motor training on the recovery of manual dexterity after primary motor cortex lesion in macaque monkeys. J. Neurophysiol. 99, 773–786. https://doi.org/10.1152/jn.01001.2007

Nudo, R.J., Milliken, G.W., 1996. Reorganization of movement representations in primary motor cortex following focal ischemic infarcts in adult squirrel monkeys. J. Neurophysiol. 75, 2144–2149.

Overduin, S.A., Zaheer, F., Bizzi, E., D’Avella, A., 2010. An instrumented glove for small primates. J. Neurosci. Methods 187, 100–104. https://doi.org/10.1016/j.jneumeth.2009.12.007

Pizzimenti, M.A., Darling, W.G., Rotella, D.L., McNeal, D.W., Herrick, J.L., Ge, J., Stilwell-Morecraft, K.S., Morecraft, R.J., 2007. Measurement of Reaching Kinematics and Prehensile Dexterity in Nonhuman Primates. J. Neurophysiol. 98, 1015–1029. https://doi.org/10.1152/jn.00354.2007

Ramanathan, D.S., Guo, L., Gulati, T., Davidson, G., Hishinuma, A.K., Won, S.-J., Knight, R.T., Chang, E.F., Swanson, R.A., Ganguly, K., 2018. Low-frequency cortical activity is a neuromodulatory target that tracks recovery after stroke. Nat. Med. https://doi.org/10.1038/s41591-018-0058-y

Rouse, A.G., Schieber, M.H., 2016. Spatiotemporal Distribution of Location and Object Effects in Primary Motor Cortex Neurons during Reach-to-Grasp. J. Neurosci. 36, 10640–10653. https://doi.org/10.1523/JNEUROSCI.1716-16.2016

Rouse, A.G., Schieber, M.H., 2015. Spatiotemporal distribution of location and object effects in reach-to-grasp kinematics. J. Neurophysiol. 114, 3268–3282. https://doi.org/10.1152/jn.00686.2015

Saleh, M., Takahashi, K., Hatsopoulos, N.G., 2012. Encoding of Coordinated Reach and Grasp Trajectories in Primary Motor Cortex. J. Neurosci. 32, 1220–1232. https://doi.org/10.1523/JNEUROSCI.2438-11.2012

Schaffelhofer, S., Scherberger, H., 2016. Object vision to hand action in macaque parietal, premotor, and motor cortices. Elife 5, e15278.

Schaffelhofer, S., Scherberger, H., 2012. A new method of accurate hand- and arm-tracking for small primates. J. Neural Eng. 9, 026025. https://doi.org/10.1088/1741-2560/9/2/026025

Schmidlin, E., Kaeser, M., Gindrat, A.-D., Savidan, J., Chatagny, P., Badoud, S., Hamadjida, A., Beaud, M.-L., Wannier, T., Belhaj-Saif, A., Rouiller, E.M., 2011. Behavioral Assessment of Manual Dexterity in Non-Human Primates. J. Vis. Exp. https://doi.org/10.3791/3258

Takahashi, K., Best, M.D., Huh, N., Brown, K.A., Tobaa, A.A., Hatsopoulos, N.G., 2017. Encoding of both reaching and grasping kinematics in dorsal and ventral premotor cortices. J. Neurosci. 1537–16. https://doi.org/10.1523/JNEUROSCI.1537-16.2016

Vaidya, M., Kording, K., Saleh, M., Takahashi, K., Hatsopoulos, N.G., 2015. Neural coordination during reach-to-grasp. J. Neurophysiol. 114, 1827–1836. https://doi.org/10.1152/jn.00349.2015

Wong, C.C., Ramanathan, D.S., Gulati, T., Won, S.J., Ganguly, K., 2015. An automated behavioral box to assess forelimb function in rats. J. Neurosci. Methods 246, 30–37. https://doi.org/10.1016/j.jneumeth.2015.03.008

